# Predicting and measuring decision rules for social recognition in a Neotropical frog

**DOI:** 10.1101/2021.08.26.457721

**Authors:** James P. Tumulty, Chloe A. Fouilloux, Johana Goyes Vallejos, Mark A. Bee

## Abstract

Many animals use signals, such as vocalizations, to recognize familiar individuals. However, animals risk making recognition mistakes because the signal properties of different individuals often overlap due to within-individual variation in signal production. To understand the relationship between signal variation and decision rules for social recognition, we studied male golden rocket frogs, which recognize the calls of territory neighbors and respond less aggressively to a neighbor’s calls than to the calls of strangers. We quantified patterns of individual variation in acoustic properties of calls and predicted optimal discrimination thresholds using a signal detection theory model of receiver utility that incorporated signal variation, the payoffs of correct and incorrect decisions, and the rates of encounters with neighbors and strangers. We then experimentally determined thresholds for discriminating between neighbors and strangers using a habituation-discrimination experiment with territorial males in the field. Males required a threshold difference between 9% and 12% to discriminate between calls differing in temporal properties; this threshold matched those predicted by a signal detection theory model under ecologically realistic assumptions of infrequent encounters with strangers and relatively costly missed detections of strangers. We demonstrate empirically that receivers group continuous variation in vocalizations into discrete social categories and show that signal detection theory can be applied to investigate evolved decision rules.

## Introduction

Animals must frequently make decisions about how to categorize natural variation in behaviorally relevant stimuli (Guilford and Dawkins 1991; Shettleworth 2010; Miller and Bee 2012). Such categorization underlies the use of communication signals for social recognition (Colgan 1983; Wiley 2013a), the phenomenon by which animals recognize individuals they have previously interacted with and assign them to social categories (e.g., “neighbor”, “offspring”, or “nest-mate”). For example, the use of vocalizations to recognize familiar individuals has been demonstrated in a variety of vertebrate taxa, including birds (Brooks and Falls 1975a; Aubin and Jouventin 2002), mammals (Cheney and Seyfarth 1980; Proops et al. 2009), amphibians (Davis 1987; Chuang et al. 2017), and fish (Myberg and Riggio 1985). Social recognition allows animals to direct behaviors to appropriate individuals but is subject to uncertainty because signal properties, such as the acoustic properties of vocalizations, often vary both among and within individuals (Gerhardt 1991). For recognition to be possible based on such signals, signal properties must be individually distinctive, meaning they vary less within individuals than among individuals (Beecher 1989). Receivers must then group within-individual variation in signal properties into discrete social categories and possess decision rules for behaviorally discriminating between signals produced by individuals belonging to these different categories. Despite many examples of social recognition (Tibbetts and Dale 2007; Wiley 2013a; Carlson et al. 2020), we still know little about how animals categorize variation in signal properties to accomplish this task (Miller and Bee 2012; Yorzinski 2017; Tumulty and Sheehan 2020).

Because communication-based social recognition is a product of coevolution between signalers and receivers, a receiver’s decision rules should reflect the natural variation in signals that occurs within and among individuals (Nelson 1989; Nelson and Marler 1990; Bee and Gerhardt 2001a, 2001b, 2001c; Aubin and Jouventin 2002; Bee 2004). However, the signal properties of different individuals often overlap, and animals aiming to discriminate between individuals based on these signal properties risk making mistakes. Additionally, the outcomes of correct and incorrect decisions may have different payoffs (in terms of fitness), and animals may differentially incur these payoffs based on the rates at which they encounter different individuals. These three factors – (1) individual variation in signals, (2) payoffs of correct and incorrect decisions, and (3) encounter rates – have likely shaped decision rules through evolution by natural selection and are thus fundamental considerations in applying signal detection theory to investigate the evolution of social recognition (Reeve 1989; Wiley 1994, 2006, 2013b). Signal detection theory and the associated acceptance threshold model (Reeve 1989) have been influential in interpreting adaptive decision rules in animals (e.g., Getty 1995; Lynn et al. 2005; Hauber et al. 2006; Steiger and Müller 2010; Mora-Kepfer 2014; Hanley et al. 2019; Rossi et al. 2019). However, studies typically rely on assumptions about the underlying distributions of cue or signal variation rather than measuring them directly to make precise predictions about decision rules.

To investigate the relationship between signal variation and decision rules for social recognition, we analyzed call variation and measured decision rules for discriminating between territory neighbors and strangers in golden rocket frogs, *Anomaloglossus beebei*. Male golden rocket frogs defend territories in bromeliads where they vocalize with pulsatile advertisement calls to attract females and advertise territory ownership (Fig. 1a, b). Territorial males recognize the advertisement calls of familiar neighbors and respond less aggressively to the calls of neighbors than to the calls of strangers (Bourne et al. 2001; Tumulty and Bee 2021), producing a phenomenon called the “dear enemy effect” (Wilson 1975; Tumulty 2018). This behavior should allow males to reduce the costs of contests (Dyson et al. 2013) that would otherwise be associated with repeated aggressive interactions with established neighbors (Getty 1987; Temeles 1994). The evolution of neighbor recognition and the dear enemy effect in golden rocket frogs has not been associated with an increase in the identity information in signals; the advertisement calls of golden rocket frogs have similar patterns of among- and within-individual variation as those of a close relative (*Anomaloglossus kaiei*) that does not discriminate between the calls of neighbors and strangers (Tumulty et al. 2021). Instead, selection has acted on receivers in golden rocket frogs, likely by modifying pre-existing mechanisms of plasticity in aggression to create decision rules that are specific to the calls of individual neighbors (Tumulty et al. 2021). The decision rules that males use to categorize calls as belonging to neighbors or strangers are currently unknown. In essence, a territorial male must decide whether an aggressive response to the call of a nearby male is warranted based on whether the call is sufficiently different from their memory of the acoustic properties of a neighbor’s calls. Such thresholds for discrimination place boundaries around perceptual categories and are sometimes referred to as “just-meaningful differences” (Nelson 1988; Nelson and Marler 1990; Gerhardt 1992). In contrast to “just-noticeable differences,” which describe the smallest perceptible differences, just-meaningful differences refer to the smallest ecologically meaningful difference in a signal property, and they can be empirically measured in playback studies as the smallest difference in a property that elicits a differential response from a receiver. Although the signal properties mediating recognition are unknown, temporal properties related to the call’s pulsatile structure are the most reliable for statistically discriminating between individuals based on calls (Pettitt et al. 2013; Tumulty et al. 2021).

**Figure 1.**
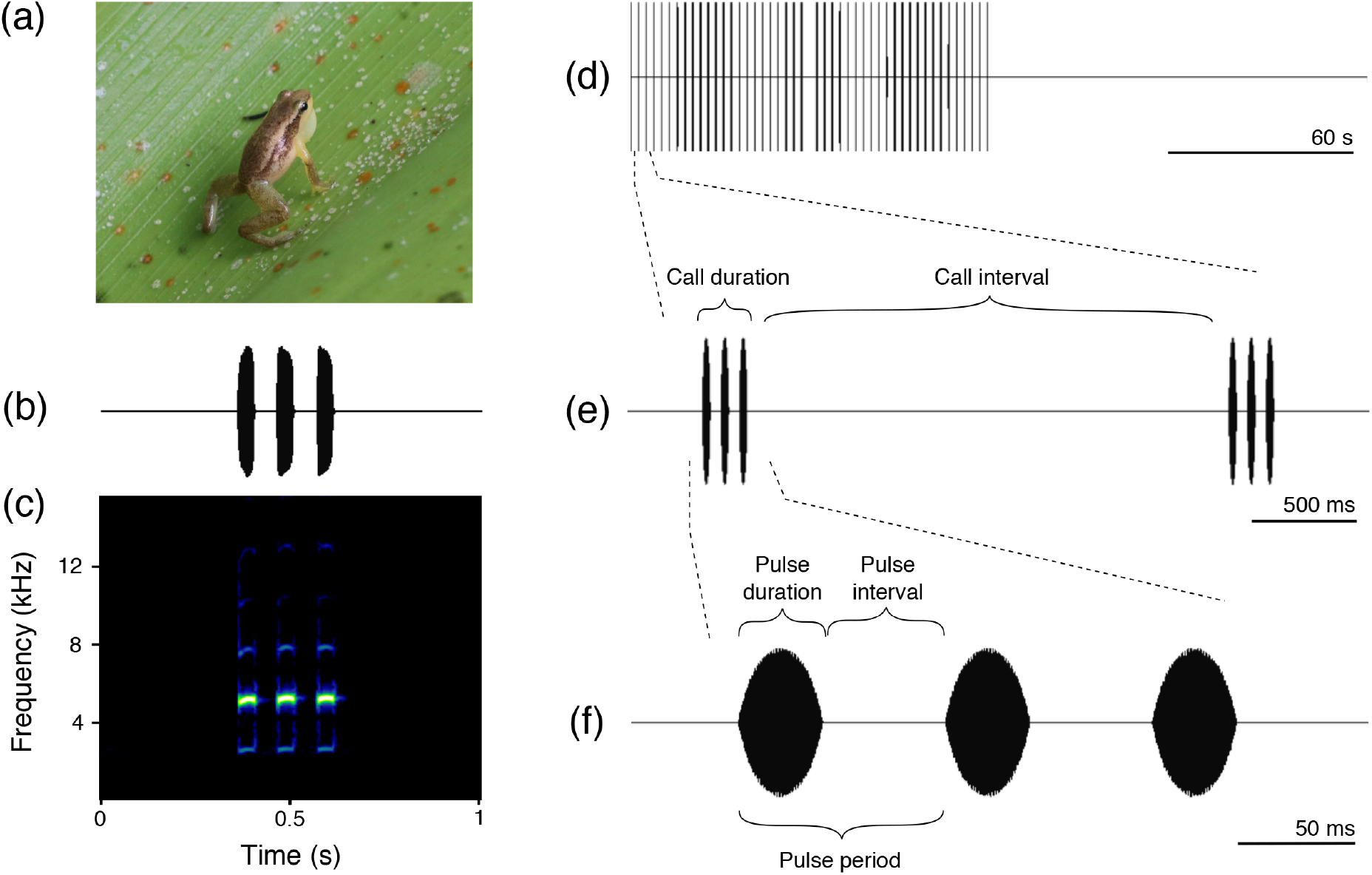
Natural and synthetic vocalizations of male golden rocket frogs. (a) Male golden rocket frogs are small (16-18mm) poison frogs that call and defend territories in large terrestrial bromeliads. (b) Waveform (amplitude over time) and (c) spectrogram (frequency over time) of a three-pulse advertisement call produced by a male golden rocket frog. (d-f) Waveforms of a synthetic advertisement call stimulus used in habituation-discrimination playbacks. (d) The full 4-minute stimulus period consisted of 2 minutes of advertisement calls followed by 2 minutes of silence. (e) A close-up of two consecutive advertisement calls showing the temporal properties of calls, and (f) a close-up of one advertisement call showing the three pulse temporal properties that were manipulated in our experiment.

This study had three objectives. First, we generated predictions about the optimal thresholds for discriminating between neighbors and strangers based on patterns of individual variation in the acoustic properties of advertisement calls, focusing on the most individually distinctive properties of pulse temporal structure. To do so, we analyzed acoustic variation in advertisement calls and used a signal detection theory model of receiver utility to predict optimal thresholds based on call variation. Second, we measured the discrimination thresholds of territorial males using a field playback experiment based on the habituation-discrimination paradigm (Rankin et al. 2009; Shettleworth 2010). Habituation is a widespread form of learning that underlies plasticity of aggression in frogs (Brenowitz and Rose 1994; Owen and Perrill 1998; Bee and Gerhardt 2001c, 2001b, 2002; Marshall et al. 2003; Humfeld et al. 2009; Tumulty et al. 2021) and has been suggested as a learning mechanism underlying neighbor-stranger discrimination in territorial species (Peeke and Peeke 1973; Peeke 1984; Petrinovich 1984). Males initially responded aggressively to calls simulating the arrival of a new neighbor by approaching the speaker and producing aggressive calls. We repeatedly played the stimulus until males met pre-determined habituation criteria indicating they were no longer responding aggressively to playbacks. Following habituation, we changed the temporal properties of pulses within stimulus calls by magnitudes spanning the range of values predicted by our signal detection theory model to empirically estimate the threshold change required to elicit a recovery of habituated aggression. A recovery of aggression would indicate that a territorial male behaviorally discriminated between the habituation and discrimination stimuli based on the experimentally manipulated properties and that it treated the discrimination stimulus as if it were a call produced by a stranger. Finally, we used the signal detection theory model to explore the combinations of stranger encounter rates and payoffs for correct and incorrect decisions that would produce the observed discrimination thresholds from our playback experiment.

## Methods

### Study system

Golden rocket frogs are a species of poison frog (Aromabatidae, Dendrobatoidea) endemic to Guyana, South America. Playback experiments were performed from June-July 2017 in Kaieteur National Park, Guyana, where these frogs live and breed in large terrestrial bromeliads (*Brocchinia micrantha*) that grow along the plateau near Kaieteur Falls. Males aggressively defend stable territories that consist of one or more bromeliads (Tumulty and Bee 2021). These plants contain reproductive resources in the form of phytotelmata, which are pools of water that collect in leaf axils and serve as oviposition and tadpole deposition sites (Bourne et al. 2001; Pettitt et al. 2018; Tumulty and Bee 2021). Males care for eggs and tadpoles in their territories, attending egg clutches and transporting tadpoles between phytotelmata (Bourne et al. 2001; Pettitt et al. 2020). Golden rocket frogs are diurnal, and males regularly vocalize from their territories in the morning with conspicuous advertisement calls. These calls consist of 1-6 short pulses (typically 2-4 pulses) with dominant frequencies between 4.6 and 5.8 kHz (Fig. 1b, c) and are typically delivered at rates of about 24 calls/min (Bourne et al. 2001; Pettitt et al. 2012; Narins et al. 2018). Territorial males respond aggressively to playbacks of conspecific advertisement calls with aggressive calls and phonotaxis (Bourne et al. 2001; Tumulty and Bee 2021).

### Acoustic recordings and measurements

We recorded advertisement calls of territorial males from a distance of 0.5-1 m using a microphone (Sennheiser ME-66, Wedemark, Germany) and digital recorder (Marantz PMD-620, Kanagawa, Japan; 44.1 kHz sampling rate, 16-bit resolution). After recording, we measured the air temperature at the male’s calling site. Some acoustic properties of frog calls can vary with temperature (reviewed in Gerhardt and Huber 2002), but there was very little temperature variation during the times when this species regularly calls (range = 22.8 to 25.8°C for our recordings). We measured the temporal and spectral properties of 20 advertisement calls from each of 20 males (400 calls total) using Raven Pro 1.5 (Cornell Lab of Ornithology, Ithaca, NY). Acoustic properties of each pulse in a call were measured separately. We measured call duration as the time from the onset of the first pulse to the offset of the last pulse in a call, and the call interval as the time from the offset of the last pulse in a call to the onset of the first pulse in the subsequent call. Call period was the sum of call duration and call interval. For each pulse in a call, we measured pulse duration as the time between the onset and the offset of the pulse and pulse interval as the time between the offset of the pulse and the onset of the subsequent pulse. The sum of pulse duration and pulse interval was pulse period. Additionally, we measured two temporal properties to describe pulse shape: pulse rise time (onset to maximum amplitude) and pulse fall time (maximum amplitude to offset). We measured the dominant frequency of each pulse using the max frequency function on the power spectrum of the pulse (1024 point, Hamming window).

### Predicting optimal decision rules

To predict optimal thresholds for discriminating between neighbors and strangers, we applied the logic of signal detection theory to the dear enemy effect. In doing so, we asked what threshold would be most useful for a territory owner aiming to detect the “signal” of a stranger’s calls while ignoring the “noise” of a neighbor’s calls. A territory owner must first form a memory of the acoustic properties of a neighbor’s calls as a result of repeatedly hearing the neighbor vocalize. This memory must be broad enough to accommodate natural levels of within-individual variation in calls. Then, upon hearing a new call, the territory owner must decide if the call is sufficiently different from its memory of its neighbor’s calls that the call is more likely to have been produced by a stranger than by a neighbor. Thus, the “noise” that males must ignore can be represented as the distribution of within-individual differences that occur in advertisement calls, while the “signal” they must detect can be represented by the distribution of among-individual differences. The territory owner must employ a decision rule for determining whether any difference between a call it just heard and its memory of a neighbor’s call is more likely due to a within-individual difference that can be ignored or to an among-individual difference that warrants an aggressive response.

We generated distributions of within-individual and among-individual differences in acoustic properties using our data set of measurements of call properties. We restricted our analysis to the first three pulses in each call. This decision allowed us to include all 20 individuals in all analyses, as six males never produced calls with more than three pulses. Indeed, in golden rocket frogs, the majority of advertisement calls consist of either two or three pulses (73% of calls in our data set) and calls with more than four pulses are exceedingly rare (Bourne et al. 2001; Pettitt et al. 2012, 2020). While the number of pulses per call can be variable, even within bouts of calling, the acoustic properties of different pulses within calls are generally highly correlated with each other (Fig. S-1). Moreover, Pettitt et al. (2012) reported that analyses of even a single pulse captured magnitudes of within-individual and among-individual differences equivalent to those based on analyses of three pulses. Thus, we could be confident that the first three pulses in calls would capture the meaningful within- and among-individual variation in calls. Prior to analysis we removed any temperature-dependent variation in acoustic properties by using linear regression (Platz and Forester 1988) to standardize values of all call properties to the mean temperature at which recordings were made (24 °C). Such standardization is common in studies of frog vocal behavior (Gerhardt and Huber 2002) and allowed us to remove any variation that was due to a known environmental influence on frog calls and not due to the individual (phenotypic) differences of interest. Analyses based on acoustic properties without temperature correction yielded very similar results and can be found in the supplementary material (Tables S-1, S-2, S-3, S-4). For each temperature-standardized property, we computed pairwise differences (as a percentage of the smaller value) between every call and every other call in the data set (Bee 2004). These pairwise differences were classified as either within-individual differences (differences between each call and the 19 other calls recorded from the same individual) or among-individual differences (differences between each call from an individual and the 380 calls recorded from the 19 other individuals in the data set). We represent these data as probability density functions of within-individual and among-individual differences for each property (Fig. 2; Fig. S-2, S-3, S-4, S-5).

**Figure 2.**
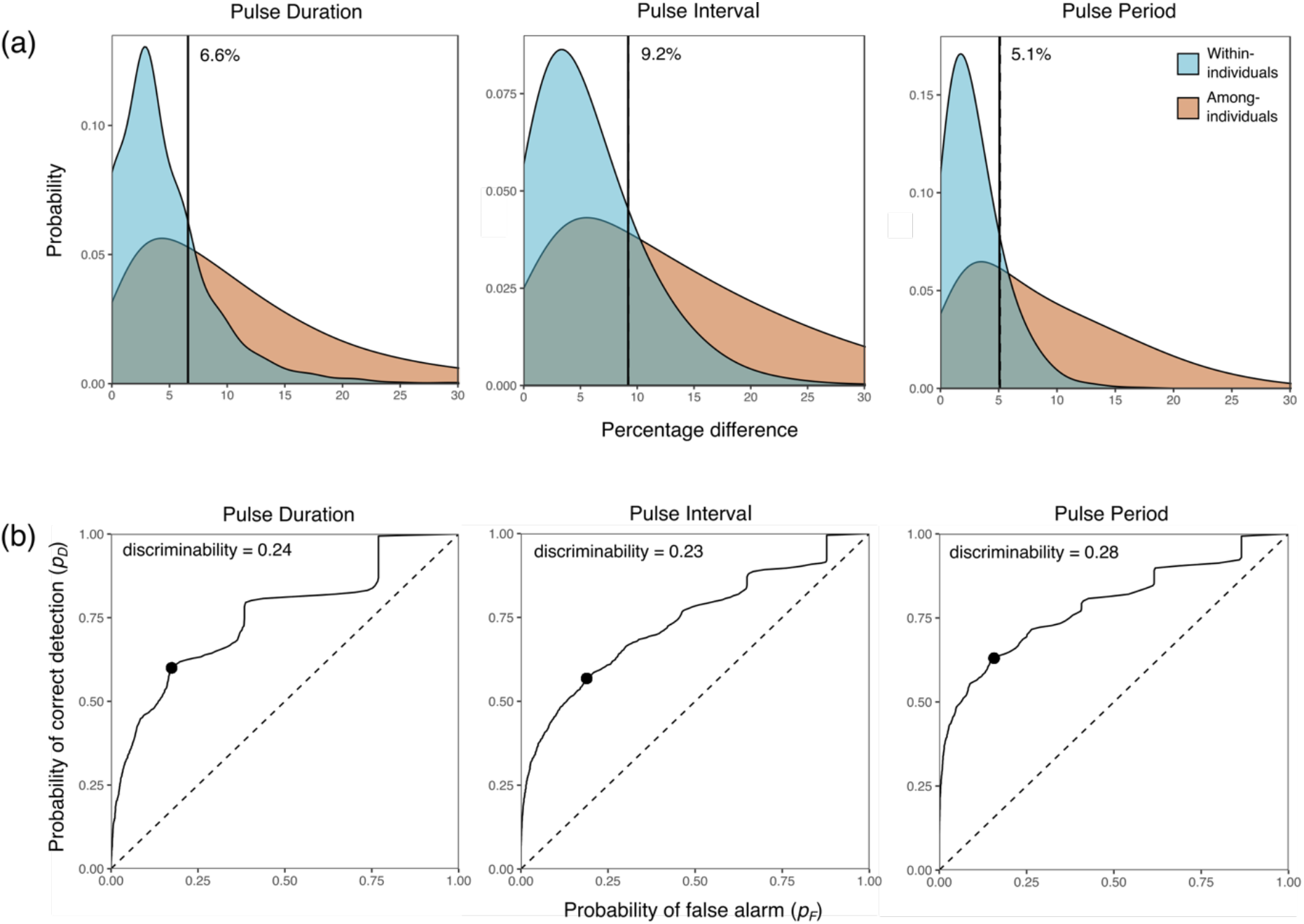
(a) Probability density functions of the within-individual and among-individual percentage differences in pulse duration, pulse interval, and pulse period. Data are pooled from all three pulses in a call. The vertical line in each graph represents the predicted optimal discrimination threshold for that call property based on signal variation, assuming equivalent neighbor and stranger encounter rates (i.e., *p_s_* = 0.5) and relative payoffs (i.e., (*r* - *f*)/(*d* - *m*) = 1)). Optimal thresholds, computed as the threshold that maximizes receiver utility, represent the percentage change in each property that was predicted to elicit behavioral discrimination from territorial males in field playback tests. (b) Receiver operating characteristic (ROC) curves generated from within-individual and among-individual differences in each acoustic property. These curves are generated by computing the probability of correct detection (*p_D_*) and probability of false alarm (*p_F_*) for each possible threshold. Points show the location of *p_D_* and *p_F_* that correspond to the predicted optimal thresholds in (a). Discriminability values were computed as the area between the ROC curve and the positive diagonal of the unit square (dotted line).

In the context of the dear enemy effect, responding aggressively to a stranger is a “correct detection,” whereas mistakenly responding aggressively to a neighbor is a “false alarm.” Correctly withholding aggression from a neighbor is a “correct rejection,” but mistakenly withholding aggression from a stranger is a “missed detection.” For each call property, and over a range of candidate discrimination thresholds (in 0.1% increments), we computed the probability of false alarm as the proportion of within-individual differences above a candidate threshold and the probability of correct detection as the proportion of among-individual differences above the same candidate threshold. These probabilities were used to create receiver operating characteristic (ROC) curves, which visualize the separation of the probability density functions representing within-individual and among-individual differences by showing the relationship between the probabilities of false alarm and correct detection over the range of possible thresholds. We further quantified this separation by computing the discriminability of among-individual differences as the area between the ROC curve and the positive diagonal of the unit square (Wiley 2006). This measure of discriminability is used in place of the traditional *d*’ in signal detection theory, which relies on assumptions of normality and equality of variance. Discriminability varies from 0, indicating completely overlapping probability density functions, to 0.5, indicating no overlap between probability density functions.

The complementary probabilities of false alarm and correct detection are the probabilities of correct rejection and missed detection, respectively. We incorporated the above probabilities in a signal detection theory model of receiver utility (Wiley 1994, 2013b) and found optimal thresholds as those that maximized utility (*U*), defined as

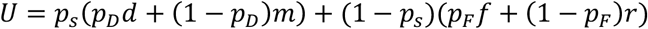

where *p_s_* = probability of encountering a “signal” in a window of time (i.e., stranger encounter rate) and 1 - *p_s_* = probability of encountering “noise” (i.e., neighbor encounter rate). Given that a stranger is calling, *p_D_* = probability of correct detection (responding aggressively), and 1 - *p_D_* = probability of missed detection (withholding aggression). Given that a neighbor is calling, *p_F_* = probability of false alarm (responding aggressively), and 1 - *p_F_* = probability of correct rejection (withholding aggression). This model also includes the payoffs (in terms of relative fitness) of correct and incorrect decisions. The payoff for a correct rejection (*r*) is the presumed best-case scenario in the context of a dear enemy effect – because it entails correctly ignoring a non-threatening neighbor – and other payoffs are considered relative to it. The payoff for correct detection (*d*) is lower because a territory owner repels a stranger but may risk injury and energy expenditure if they need to fight the stranger to do so (e.g., Dyson et al. 2013). The payoff for a false alarm (*f*) is lower still because a neighbor that has been wrongly attacked should be more likely to escalate a fight than a stranger because it has already invested in establishing and defending its territory, whereas the stranger lacks anything to defend (e.g., prior residence effects; (Davies 1978; Baugh and Forester 1994; Yang et al. 2020). Finally, the payoff for missed detection (*m*) of a stranger is considered the worst-case scenario because an un-detected stranger could, for example, steal a mating with a female, make use of a defended resource, or take over an entire territory (e.g., relative threat hypothesis; Getty 1987; Temeles 1994; Booksmythe et al. 2010; Siracusa et al. 2017). While we do not have any measurements of these payoffs in nature, we generally expect that in the dear enemy context, *r* > *d* > *f* >> *m*. These payoffs influence optimal thresholds based on the ratio of differences between correct and incorrect responses to neighbors (*r* - *f*) and strangers (*d* - *m*), that is, (*r* - *f*)/(*d* - *m*) (see Wiley 1994). Given that a neighbor is calling, the difference between the payoffs of correct rejection and false alarm (*r* - *f*) is the cost of a false alarm. Given that a stranger is calling, the difference between the payoffs of correct detection and missed detection (*d* - *m*) is the cost of a missed detection. Note that by defining optimal thresholds as those that maximize the utility equation above, we are implicitly assuming that encounter rates are fixed and uniform.

We first present predictions based on assumptions that neighbor and stranger encounter rates are equal (*p_s_* = 0.5) and the payoff differences between correct and incorrect responses to neighbors and strangers are equal ((*r* - *f*)/(*d* - *m*) = 1). This combination of assumptions predicts optimal thresholds equivalent to those predicted by a method that simply maximizes the difference between the probability of correct detection (*p_D_*) and the probability of false alarm (*p_F_*) and, therefore, yields predictions based only on patterns of signal variation. These initial threshold predictions thus serve as a null model of the effect that encounter rates and payoffs have in shifting optimal thresholds; if observed thresholds deviate from our initial predictions, we can then use the observed thresholds to infer how encounter rates and payoffs may have shifted the optimal threshold.

We focused our treatment of predicted optimal thresholds on the temporal properties of pulse duration, interval, and period. The predicted optimal thresholds for other call properties can be found in the supplementary material. Additionally, the same acoustic properties measured from different pulses within a call were treated as separate variables when computing pairwise differences, but we pooled these differences together when generating within-individual and among-individual distributions and predicting thresholds because the distributions for different pulses were always similar. Distributions and predicted optimal thresholds for each pulse analyzed separately can be found in the supplementary material.

### Measuring decision rules

#### (i) Playback stimuli

We used a custom sound synthesis program (written by J. J. Schwartz) to generate synthetic calls (Fig. 1 d, e, f) that were modeled after natural calls but that allowed us to systematically manipulate acoustic properties during habituation-discrimination playbacks. Individual pulses consisted of three harmonically related, phase-locked sinusoids with frequencies (and relative amplitudes) of 2669 Hz (−30 dB), 5338 Hz (0 dB), and 8007 Hz (−30 dB), such that 5338 Hz was the dominant frequency. We edited pulses into calls using Adobe Audition 1.5 (Adobe Systems Inc., San Jose, CA, USA). A call for the habituation stimulus represented the population mean for all properties and consisted of three 36-ms pulses separated by 52-ms intervals, resulting in a pulse period of 88 ms (Fig. 1f). Each pulse was shaped with 16-ms exponential convex onset and offset ramps that reached 50% maximum amplitude at 4 ms after onset and before offset, respectively. We created a 1-minute-long stimulus consisting of 23 calls at a rate of 1 call/2.5 s. We appended two repetitions of this stimulus followed by two minutes of silence to create a 4-minute stimulus period (Fig. 1d). These stimuli were saved as WAV-files (44.1 kHz sampling rate, 16-bit resolution). During the discrimination phase of the experiment, we presented a novel stimulus that was designed to simulate a different individual based on differences in the temporal properties of pulses. To estimate a behavioral threshold for responding aggressively following habituation, we tested four novel stimuli that featured a 3%, 6%, 9%, or 12% change in pulse temporal properties. These values were selected to span the range of optimal discrimination thresholds predicted by our signal detection theory model under the assumptions of equal encounter rates and payoffs for interactions with neighbors and strangers (see below). These manipulations involved changing three inter-related temporal properties of pulses (pulse duration + pulse interval = pulse period) simultaneously and by the same percentage difference to account for their natural intercorrelation. All other acoustic properties were held constant between stimuli used during the habituation and discrimination phases. For each experimental treatment, half of the males heard an increase in pulse temporal properties, and half heard a decrease. As a control, we also included a 0% change treatment, which consisted of additional broadcasts of the habituation stimulus during the discrimination phase. Stimuli for all five treatments had the same pulse duty cycle (pulse duration/pulse period) of 0.41.

#### (ii) Playback protocol

We performed 62 habituation-discrimination playback tests with 17 male subjects. This species is sexually dichromatic, and males are easily identified in the field (Engelbrecht-Wiggans and Tumulty 2019). Additionally, males were photographed so that they could be individually identified based on individually distinctive dorsal patterns. All playbacks began in the morning (0700 – 1100 hours), when this species is most active. Stimuli were played from a digital audio player (iPod, Apple, Cupertino, CA, USA) connected to an amplified field speaker (Saul Mineroff Electronics, Elmont, NY, USA). The speaker was calibrated to produce stimuli at 80 dB SPL measured at 1 m, which is within the range of variation in call amplitude for this species (Pettitt et al. 2012). To begin a test, we chose a calling male and positioned the speaker on a tripod 1.5 – 2 m from the subject in a direction with no adjacent neighbors. The speaker was positioned at an equivalent height to the subject and so that it contacted a bromeliad leaf to simulate a realistic calling location. We also set up a microphone (Sennheiser ME-66, Wedemark, Germany) and digital recorder (Marantz PMD-620, Kanagawa, Japan; 44.1kHz sampling rate, 16-bit resolution) to record the subject’s vocal responses. Subjects sometimes stopped calling during speaker set-up, so we waited at least 5 minutes or until it resumed calling to begin the playback.

We started a test with a 4-minute pre-stimulus period, during which we observed the subject’s behavior in the absence of any stimulus. We then began the habituation phase, during which we repeatedly played the 4-min habituation stimulus (Fig. 1d). This phase of the test simulated the arrival of a new neighbor. During each minute of the test, we recorded four behavioral measures of aggression: (1) the number of aggressive calls, (2) the number of pseudo-aggressive calls, (3) the subject’s approach distance, and (4) the subject’s closest position to the speaker. Golden rocket frogs have distinct aggressive calls (Bourne et al. 2001; Pettitt et al. 2012), which consist of long trains of pulses typically preceded by several introductory pulses with relatively longer inter-pulse intervals. We classified all calls with at least seven pulses as aggressive calls. Calls consisting of the characteristic introductory pulses without a subsequent train of pulses were scored as “pseudo-aggressive calls” (Tumulty et al. 2021). Call counts were always confirmed from the audio recordings of each test. We also noted the subject’s position at the start of each minute of the test as well as its closest position to the speaker during each minute. After the test, we measured the distances from the speaker to each noted position, and from these measurements, we computed the subject’s approach distance (net displacement towards the speaker) during each minute as well as the distance from the speaker to the subject at its closest position to the speaker during each minute. We also recorded air temperature at the final location of the frog. The range in temperatures during our playbacks was 22.4 to 26.8 °C (mean = 24.3 °C).

We aborted the test if the subject did not show any aggressive behaviors during the first three stimulus periods of the habituation phase (i.e., 12 min), started interacting aggressively with a real male, or started courting a female. We played the habituation stimulus for at least six stimulus periods (i.e., 24 mins) and until the subject met our habituation criteria of no aggressive calls, pseudo-aggressive calls, or approach movements towards the speaker for three consecutive stimulus periods (maximum habituation time was 5.6 hrs). Once these habituation criteria were met, we switched to the designated discrimination stimulus (i.e., a 0% [control], 3%, 6%, 9%, or 12% change) for three stimulus periods during the discrimination phase. After the test, we caught the male, if possible, to confirm its identity. While we did not always succeed in catching subject males after tests, their territories are remarkably stable (Tumulty and Bee 2021), and every time we caught a male it was the same individual as assumed based on location. We attempted to test each subject in all five treatments; however, some tests were unsuccessful for a variety of reasons (see above), so not all 17 subjects were tested in all treatments. We allowed at least five days between consecutive tests to minimize the potential effects of long-term habituation to the habituation stimulus. There was no effect of number of attempted playback tests on a subject’s likelihood of response (logistic mixed effects model: *χ*^2^ = 2.4, *p* = 0.12, *n* = 118 attempted tests), nor was there an effect on their time to reach habituation criteria (linear mixed effects model: *χ*^2^ = 0.2, *p* = 0.65, *n* = 62 successful tests), one measure of overall aggressiveness. The final sample sizes for each treatment (and sample sizes for the corresponding increases and decreases in pulse properties) were as follows: 0%, *n* = 12; 3%, *n* = 12 (6 increased and 6 decreased, respectively); 6%, *n* = 12 (6 and 6); 9%, *n* = 13 (6 and 7); and 12%, *n* = 13 (6 and 7). Treatment order was randomly assigned to each subject, and we changed the file names of the stimuli so that observers were blind to treatments when conducting playbacks and scoring aggressive behaviors.

#### (iii) Statistical analyses

To examine the effects of treatments on aggressive responses, we summarized data by block, which consisted of three consecutive stimulus periods. For each block, we summed the number of aggressive and pseudo-aggressive calls and summed the approach distances. To control for slight variation between tests in initial speaker distance, we expressed the subject’s closest position to the speaker as a proportion of the total distance between the speaker and subject at the start of the test; we used the inverse of this proportion so that higher values indicate a closer approach to the speaker. We subjected these four correlated measures of aggression to a principal components transformation (centered to zero and scaled to unit variance) using the ‘prcomp’ function in R (R Core Team 2017) and used the first principal component as an aggression index. This aggression index explained 56% of the overall variation in aggressive behaviors and was positively correlated with the number of aggressive calls (loading = 0.52) and pseudo-aggressive calls (0.40), approach distance (0.57), and closest position to the speaker (0.50). Additionally, results for each measure of aggression are plotted separately in the supplementary material (Fig. S-6, S-7, S-8, S-9). To estimate a behavioral threshold for behaviorally discriminating between neighbors and strangers, we tested for a change in aggression index between the last block of the habituation phase and the single block of the discrimination phase using Wilcoxon signed rank tests within each treatment. For these tests we report *V* statistics, which correspond to the sum of positive ranks. We estimated the behavioral threshold as occurring in the range between the smallest percentage change in the novel stimulus that elicited significant recovery of aggression and the largest change that did not.

### Modeling encounter rates and payoffs

After estimating the threshold for behaviorally discriminating between neighbors and strangers in the playback tests, we returned to our signal detection theory model and explored the ranges of parameter values for stranger encounter rates and relative payoffs that could result in the observed behavioral threshold. Because classic methods for determining parameters based on measured thresholds rely on assumptions of normality and equal variance (Gescheider 1976), which our data did not meet, we instead determined parameter values by computing optimal thresholds from the receiver utility equation for a large range of possible values of stranger encounter rates and relative payoffs. We varied stranger encounter rates (*p_s_*) from 0 to 1, in 0.1 increments. To vary relative payoffs ((*r* - *f*)/(*d* - *m*)), we varied values of r and d such that the resulting numerator and denominator of this ratio each varied from 1 to 20, allowing us to explore parameter space ranging from equivalent costs of false alarms (*r* - *f*) and missed detections (*d* - *m*) (i.e., (*r* - *f*)/(*d* - *m*) = 1) to scenarios in which the costs of false alarms are twenty times greater than the costs of missed detections ((*r* - *f*)/(*d* - *m*) = 20) and scenarios in which the costs of missed detections are twenty times greater than the costs of false alarms ((*r* - *f*)/(*d* - *m*) = 0.05). We then selected the combinations of these parameters that resulted in the optimal thresholds estimated from our playback tests.

## Results

### Predicted optimal decision rules

As illustrated in Figure 2a (see also Fig. S-2, S-3, S-4, S-5), there was overlap in the distributions of within-individual and among-individual differences in acoustic properties, thus demonstrating that these properties vary continuously within the population. Importantly, however, among-individual differences were larger than within-individual differences for all acoustic properties. Compared with other properties, the temporal properties of pulses (pulse duration, pulse interval, and pulse period) and spectral properties had larger separations between distributions representing within-individual differences and among-individual differences (Table S-1, S-3; Fig. 2a; Fig. S-2, S-3, S-4, S-5). This separation is visualized in the ROC curves in Figure 2b, which are greatly offset from the diagonal. Pulse period had a slightly larger value of discriminability (0.28) than pulse duration (0.24) and pulse interval (0.23) (Fig. 2b, Table S-1, S-3). Most within-individual differences in pulse duration, pulse interval, and pulse period were less than 5% (Fig. 2a). While small among-individual differences (<5%) in these three properties were not uncommon (constituting 25% to 36% of comparisons), most among-individual differences in pulse duration, pulse interval, and pulse period were above 5% (Fig. 2a). Assuming equivalent relative payoffs ((*r* - *f*)/(*d* - *m*) = 1) and equal neighbor and stranger encounter rates (*p_s_* = 1 -*p_s_* = 0.5), the predicted optimal thresholds for discriminating between individuals based on the temporal properties of pulses ranged between 5% and 10%: for pulse duration and pulse period the predicted optimal thresholds were 5.1% and 6.6%, respectively, while that for pulse interval was somewhat higher at 9.2% (Fig. 2a, Table S-1). The distributions of within-individual and among-individual differences and predicted thresholds using data pooled across pulses (Fig. 2a, Table S-1) were very similar to those for each pulse analyzed separately (Table S-1, S-3, Fig. S-2, S-3, S-4).

### Measured decision rules

Territorial males responded to synthetic call playbacks with aggressive behaviors that were typical of responses to playbacks of natural calls (Tumulty and Bee 2021) as well as natural aggressive interactions between males (personal observations). Males often approached the speaker while the stimulus was playing and retreated to the centers of their bromeliads, away from the speaker, during intervening periods of silence. Some males approached all the way to the speaker, apparently searching for the simulated intruder. Males also responded to playbacks with aggressive calls, which were typically produced during the periods of silence between stimuli. These aggressive responses gradually decreased with repeated broadcasts of the stimulus during the habituation phase (Fig. 3), but the amount of time it took for subjects to meet habituation criteria varied considerably among individuals. The median amount of time to reach these criteria was 1 hr (or 15 stimulus periods) and ranged from 24 min (6 stimulus periods) to 5.6 hrs (84 stimulus periods).

**Figure 3.**
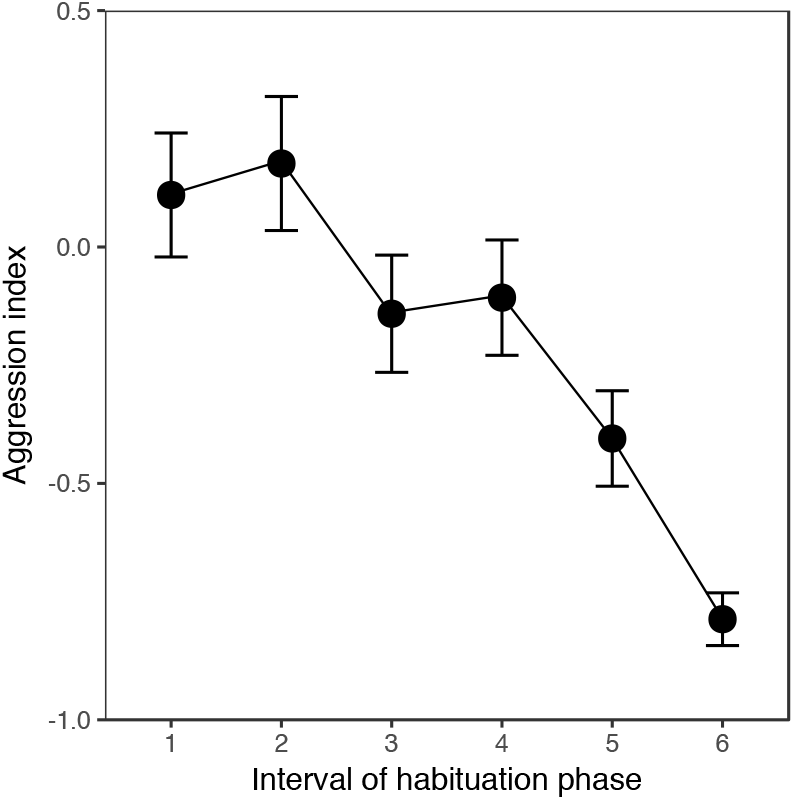
Mean (±SE) aggression index of territorial male golden rocket frogs to synthetic advertisement call playbacks over the course of the habituation phase (*n* = 62 tests of 17 males). Because subjects took different amounts of time to reach habituation criteria, here we visualized trends in aggression during the habituation phase by partitioning this phase into 6 intervals (6 being the minimum number of stimulus periods), each constituting 16.7% of the total time a subject required to reach the criteria. To do this we first summarized data in the same manner as described in the methods but by stimulus period. We then subjected these data to a principal components transformation to compute an aggression index and calculated the mean aggression index during each interval across subjects.

The threshold for behaviorally discriminating between the calls of different simulated individuals based on the percentage differences in pulse temporal properties was above 9% but below 12%. Only a 12% change in the temporal properties of pulses elicited an obvious and statistically significant recovery of aggression during the discrimination phase (*V* = 44, *p* = 0.013, *n* = 13; Fig. 4). This treatment corresponds approximately to a 10-ms change in pulse period, a 4-ms change in pulse duration, and a 6-ms change in pulse interval. In this treatment group, there was no difference in recovery of aggression between subjects that experienced an increase versus a decrease in the values of pulse temporal properties (Wilcoxon rank sum test, W = 18.5, *p* = 0.77; *n_1_* = 6, *n_2_* = 7), indicating that subjects were responding to the change in these properties instead of revealing a bias for responding with more or less aggression to longer or shorter pulse temporal properties. The median difference in the aggression index between the last block of the habituation phase and the block of the discrimination phase was zero for the 0%, 3%, 6%, and 9% treatments (Fig. 4b). A 9% change elicited recovery of aggression in some subjects (Fig. 4) during the discrimination phase, but the difference from the last block of the habituation phase was not statistically significant (*V* = 18, *p* = 0.14, *n* = 13; Fig. 4). Similarly, there were no statistically significant differences in aggression between the last block of the habituation phase and the block of the discrimination phase for the 0% change control (*V* = 9, *p* = 0.20, *n* = 12), 3% change (*V* = 16, *p* = 0.29, *n* = 12), or 6% change (*V* = 33, *p* = 0.61, *n* = 12) treatments (Fig. 4). An examination of the separate components of the aggression index suggest significant recovery of aggression in the 12% treatment was driven primarily by renewed physical approaches towards the playback speaker (Figs. S-8, S-9) and not renewed aggressive calling (Figs. S-6, S-7).

**Figure 4.**
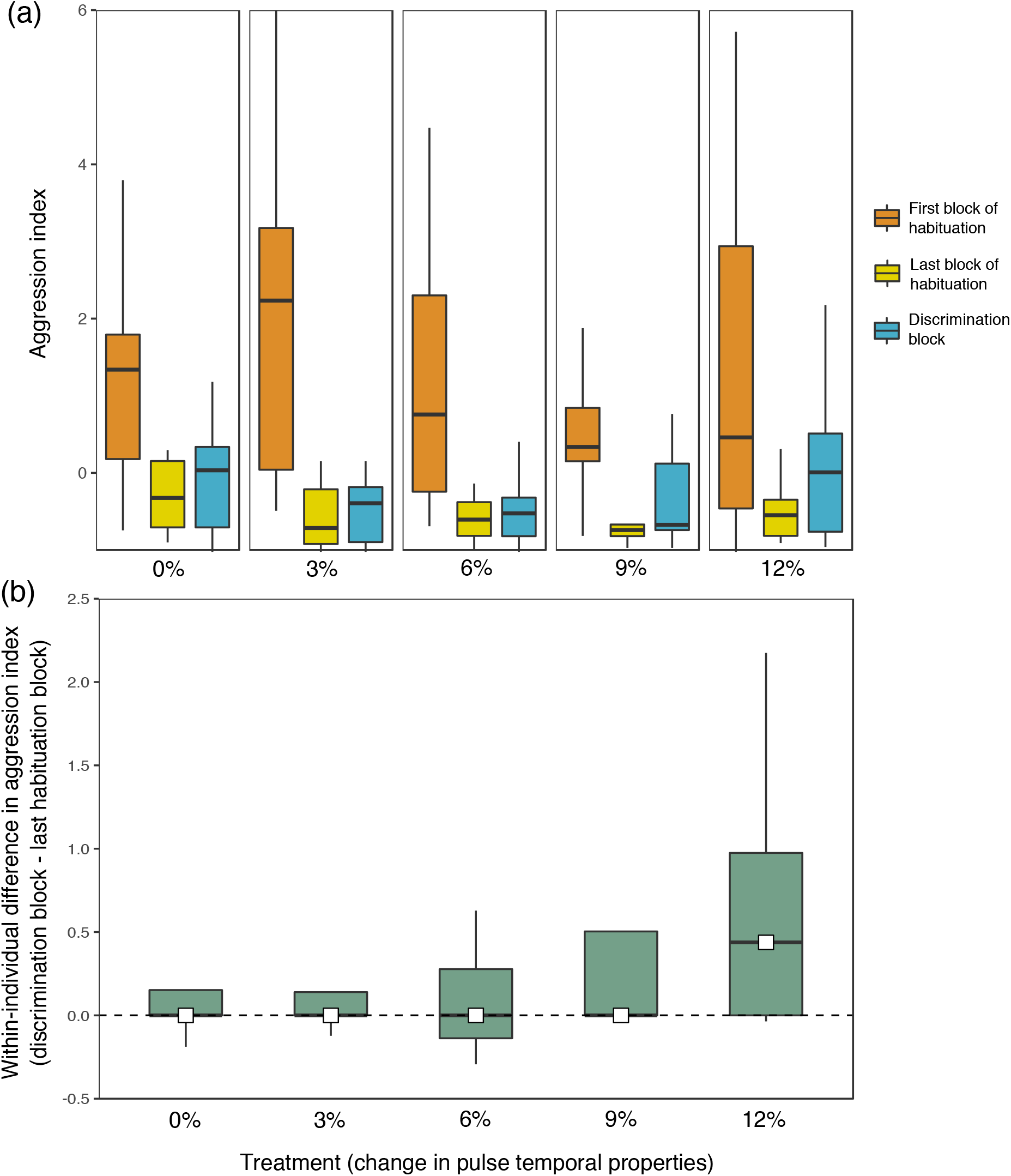
(a) Boxplots of the aggression index of territorial male subjects during the first and last blocks of the habituation phase and during the block of the discrimination phase for each treatment (0%, *n* = 12; 3%, *n* = 12; 6% *n* = 12; 9% *n* = 13; 12%, *n* = 13). (b) Boxplots showing the within-subjects difference in aggression index between the last block of the habituation phase and the discrimination phase for each treatment. Horizontal bars represent the median, box hinges represent the interquartile range, and whiskers extend to the range but no further than 1.5 times the interquartile range. White squares also show the median in (b) to aid visualization.

### Modeling encounter rates and payoffs

Exploring the range of parameters that could have produced the observed behavioral threshold of 9-12% revealed some differences from our starting assumptions of equal neighbor and stranger encounter rates (i.e.,*p_s_* = 1-*p_s_* = 0.5) and equivalent relative payoffs ((*r* - *f*)/(*d* - *m*) = 1) that produced our initial prediction of an optimal discrimination threshold between 5% and 10%. The range of possible parameter values that could have resulted in the observed threshold of 9-12% did not overlap with our initial assumptions for pulse duration and pulse period but did overlap with those assumptions for pulse interval (Fig. 5). For pulse duration and pulse period, at equivalent relative payoffs, the observed threshold could have been produced by lower stranger encounter rates. Alternatively, if stranger encounter rates are kept at 0.5 for these properties, the observe thresholds could have been produced by increased costs of false alarms (mistakenly attacking a neighbor) (Fig. 5). For all three properties, the observed thresholds are also consistent with ecologically realistic scenarios in which stranger encounter rates are rare and the costs of missed detections of strangers are greater than the costs of false alarms (lower left quadrants of Fig. 5).

**Figure 5.**
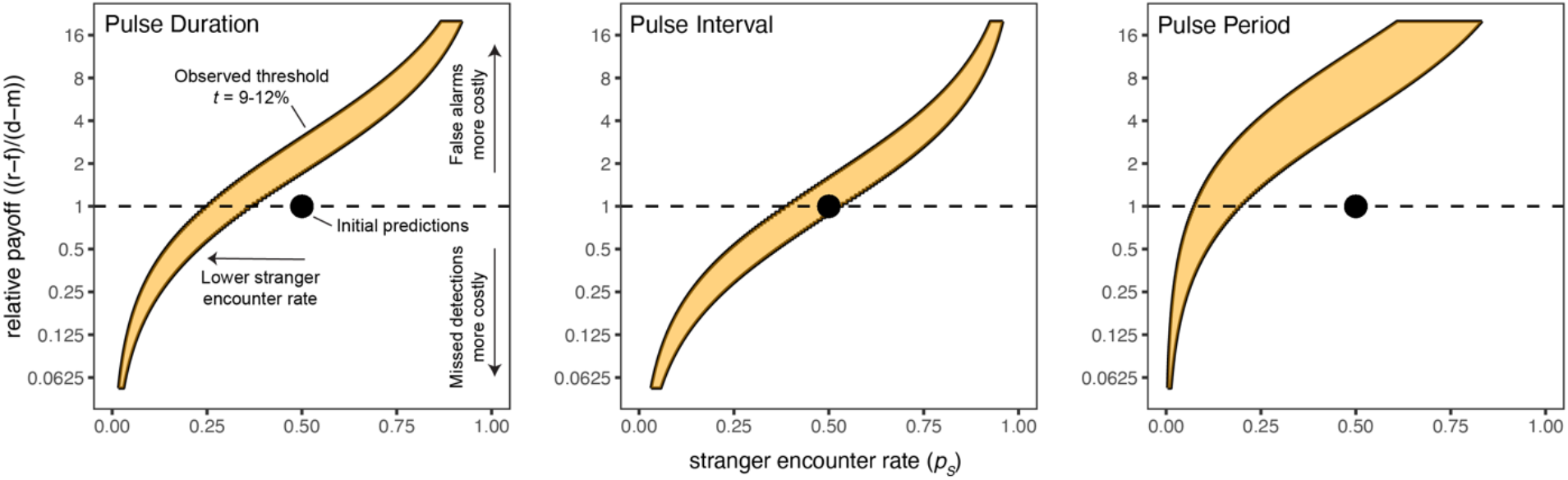
Parameter space (orange curve) for the joint values of stranger encounter rates (*p_s_*) and relative payoffs ((r-f)/(d-m)) that could result in the observed behavioral discrimination threshold of between 9% and 12% changes in pulse temporal properties. The point in each plot displays the initial assumptions of equal encounter rates to neighbors and strangers (*p_s_* = 0.5) and equivalent costs of false alarms and missed detections ((r-f)/(d-m) = 1).

## Discussion

In this study, we addressed two major questions about social recognition: what are the decision rules used to discriminate among individuals, and how are these rules shaped by patterns of individual variation in signals, the rates of encountering different individuals, and the payoffs associated with interactions with different individuals? Similar to responses to repeated playbacks of natural calls (Tumulty et al. 2021), repeated exposures to synthetic calls simulating a new neighbor led to both habituation of territorial aggression and the formation of a memory of individually distinctive signal properties. Males employed a threshold for behaviorally discriminating individual differences in calls that required a difference in pulse temporal properties between 9% and 12%: changes from the habituation stimulus of 9% or less failed to elicit recovery of aggression while a 12% change elicited renewed territorial aggression. This empirically determined range for a threshold partially overlaps, but extends somewhat higher than, the predicted range of 5% to 10% based on our initial signal detection theory model. Notably, however, our initial predictions were based on patterns of signal variation alone, as they assumed both that neighbors and strangers were encountered at equivalent rates and that the costs of false alarms (incorrectly attacking a neighbor) and missed detections (incorrectly ignoring a stranger) were equivalent. We interpret the relatively close correspondence between observed and predicted thresholds as evidence that patterns of individual variation in signals have been an important factor shaping the evolution of behavioral decision rules in the golden rocket frog’s recognition system. Nevertheless, exploring the parameter space of the model that produced thresholds in the range of 9-12% showed that both lower rates of encountering strangers than neighbors and differences in the costs of false alarms and missed detections could improve the correspondence between observed and predicted thresholds. Understanding the importance of these two additional factors in broader ecological and evolutionary contexts first requires consideration of likely encounter rates and costs of errors in natural settings.

The assumption that strangers and neighbors are encountered at equal rates is almost certainly not true for most territorial animals. Male golden rocket frogs, for example, typically call from their spatially clumped territories every day, often for several hours each day (Bourne et al. 2001; Pettitt et al. 2012, 2020; Tumulty and Bee 2021). Because of this, territorial males likely hear the calls of their nearby neighbors more frequently than they hear the calls of strangers and thus risk paying the costs of false alarms (incorrectly attacking a neighbor) more frequently than the costs of missed detection (incorrectly ignoring a stranger). Male golden rocket frogs may have evolved somewhat higher discrimination thresholds than predicted based on signal variation alone to decrease the rate at which they pay the costs of false alarms (Fig. 5). Of course, the relative payoffs themselves also impact optimal thresholds in addition to the rates at which different payoffs are incurred. Our initial predictions assumed that the payoff differences of correct and incorrect responses to neighbors and strangers were equal, an assumption that may not hold in many cases. The model showed that, at equal encounter rates, our observed threshold could have been produced by the costs of false alarms being greater than the costs of missed detections. Relatively high costs of false alarms could be the case if neighbors are particularly likely to escalate fights when challenged, even with established “dear enemies”. This scenario would be consistent with prior-residence effects in which territory holders are more likely to escalate and win contests over intruders (Davies 1978). Prior residence effects are well documented among territorial animals (Kokko et al. 2006), including in another poison frog (Baugh and Forester 1994; Yang et al. 2020). In the context of the dear enemy effect, however, the costs of missed detections may ultimately be greater than the costs of false alarms. Strangers are generally thought to pose a greater threat to territory owners than neighbors, because, unlike neighbors who already have territories of their own, strangers represent individuals who may attempt to take over another male’s defended resources (Getty 1987; Temeles 1994). A false alarm may result in a brief fight with a neighbor, which would be worse than correctly ignoring that neighbor. But the difference in payoffs between correct and incorrect responses to strangers may be greater because strangers moving through a territory network that are quickly detected can be driven off with an aggressive display, but strangers that are un-detected could potentially steal a mating with a female, make use of defended reproductive resources, or possibly usurp an entire territory. Even a stranger establishing a new territory within an existing territory network represents the addition of another nearby competitor for limited resources (Getty 1987). Relatively higher costs of missed detections than false alarms decrease the predicted optimal threshold. Overall, while we presently lack the necessary data to make reliable estimates for encounter rate and payoff parameters, the signal detection theory model fits the observed threshold under ecologically realistic conditions of low stranger encounter rates, even when failing to respond to a stranger (missed detection) is more costly than mistakenly attacking a neighbor (false alarm) (Fig. 5, lower left quadrant). Our approach of parameterizing our signal detection theory model with measures of signal variation was useful for predicting receiver decision rules in the dear enemy context and for interpreting how encounter rates and payoffs may have shaped these decision rules, either over evolutionary time or perhaps as a result of behavioral plasticity in social environments where encounter rates and costs of errors fluctuate (e.g., Stoddard 1996). We think this approach, as well as signal detection theory more broadly, will also yield useful insight when applied to other contexts of animal communication (Wiley 1994, 2006, 2013b; Scharf et al. 2020; Sumner and Sumner 2020).

An important finding from this study is the empirical determination that receivers use individually distinctive acoustic cues for discriminating between individuals. While acoustic and statistical analyses typically support the widely held view that animal vocalizations possess individually distinctive properties (Bee et al. 2016), surprisingly few previous studies have identified specific signal properties used by receivers to discriminate among individuals. Studies of the perceptual basis of the dear enemy effect in white-throated sparrows (Brooks and Falls 1975b), field sparrows (Nelson 1989) and North American bullfrogs (Bee and Gerhardt 2001a, 2001b, 2001c, 2002) have identified “pitch” (i.e., acoustic frequency) as an individually distinctive property of vocalizations that can be used to discriminate between neighbors and strangers. Our finding that male golden rocket frogs can learn to discriminate individual differences in the temporal properties of vocalizations reveals a new perceptual mechanism underlying the dear enemy effect. Across animal groups, temporal patterns in acoustic signals convey biologically important information (e.g., Pollack and Hoy 1979; Stratton and Uetz 1983; Illes et al. 2006). Among frogs, temporal information plays key roles in species recognition (Gerhardt 2001; Vélez et al. 2012), mate choice (Gerhardt et al. 1996; Akre and Ryan 2010; Tanner et al. 2017), and male-male competition (Wells and Schwartz 1984; Gerhardt et al. 2000; Burmeister et al. 2002). Our study extends the role of temporal information in frog communication to social recognition. In golden rocket frogs, the temporal properties of pulses within calls are the most individually distinctive properties of advertisement calls (Pettitt et al. 2013), and males in our experiment learned to recognize pulse temporal properties of the calls of simulated new neighbors, discriminating between calls that differed by 12% in these properties. The pattern of habituation and discrimination we observed in our experiment would allow territorial males to ignore neighbors and respond aggressively to a large subset of strangers, thereby producing a dear enemy effect based solely on patterns of individual variation in pulse temporal properties. Whether males also use other acoustic properties of calls to recognize neighbors remains to be investigated.

The magnitude of change in pulse temporal properties that was discriminated by male golden rocket frogs is noteworthy and deserves additional comment. The percentage change of 12% in the discrimination phase corresponded to absolute changes of approximately 10 ms in pulse period, 4 ms in pulse duration, and 6 ms in pulse interval. Similar sensitivity of the anuran auditory system to such small differences in temporal properties has been demonstrated previously, but only in studies of unlearned behaviors related to species recognition and mate choice conducted under highly controlled laboratory conditions (Gerhardt and Schul 1999; Bush et al. 2002). Neurophysiological studies indicate that such temporal discrimination likely involves neurons in the anuran midbrain that are tuned to particular pulse durations and intervals (Edwards et al. 2002; Rose 2014). However, that a frog can discriminate small temporal differences in acoustic signals on the order of 4 ms to 10 ms under natural conditions based on learning, as we have shown, is unprecedented in the literature and deserves additional study. At present, for example, it remains to be determined whether golden rocket frogs perceive individual differences in vocalizations categorically (Tanner and Tumulty 2020), similar to the categorical perception in songbirds that underlies discrimination between types of vocalizations (Nelson and Marler 1989) or beak colors (Caves et al. 2018). It will also be important to discover whether frogs can perceptually discriminate between smaller differences than they behaviorally discriminated in our experiment, that is, whether their just-noticeable differences are smaller than their just-meaningful differences.

We emphasize that there are many aspects of territory establishment and interactions with neighbors that were not captured in our playback experiment. Territory disputes, for example, often escalate to intense bouts of physical aggression in diverse animal species (e.g., fiddler crabs (Booksmythe et al. 2010), chameleons (Ligon 2014), cichlid fish (O’Connor et al. 2015), songbirds (Krebs 1982), and chimpanzees (Wrangham 1999)). Such interactions are also common among territorial frogs (Dyson et al. 2013), and in golden rocket frogs, physical aggression involves wrestling and chasing that can sometimes span several days (personal observations). Such physical and visual interactions create the potential for territory holders to acquire information about the reliability and fighting ability of neighbors and associate that information with memories of their neighbors’ communication signals. While these interactions are certainly important in real dear enemy relationships, the habituation-discrimination experiment nevertheless allowed us to experimentally probe the decision rules used to socially categorize variation in calls. As a taxonomically widespread form of non-associative learning, habituation likely plays a key role allowing territory holders to learn about their neighbors and adjust their aggression accordingly (Shettleworth 2010), as demonstrated in previous studies of fish (Peeke and Peeke 1973; Peeke 1984), frogs (Owen and Perrill 1998), and songbirds (Petrinovich 1984). Previous studies in other frogs, for example, have shown repeated exposure to conspecific calls can lead to habituation of aggression that can enable neighbor recognition and the dear enemy effect (Bee and Gerhardt 2001b, 2001c, 2002) or simply allow males to modulate their aggression in a way that allows them to track changes in the local density of conspecific males in the chorus (Brenowitz and Rose 1994; Marshall et al. 2003; Reichert 2010). We have recently shown that the specificity of social categories that are learned through habituation has been a key target of selection enabling a dear enemy effect in golden rocket frogs (Tumulty et al. 2021). A closely related species that does not discriminate between neighbors and strangers (Kai rocket frogs, *A. kaiei*) exhibits similar patterns of habituation to conspecific calls, but unlike golden rocket frogs, Kai rocket frogs show habituation that is generalized to conspecific calls as they do not discriminate individual differences in calls (Tumulty et al. 2021). Thus, it appears that key parameters of habituation, such as stimulus specificity, are evolutionarily labile in frogs and have been shaped by natural selection to allow individuals to navigate their social environment.

The abilities of animals to categorize continuous variation in acoustic signals underlies the recognition of mates (Gerhardt 2001; Baugh et al. 2008), rivals (Nelson 1989; Nelson and Marler 1990; Amézquita et al. 2011), and offspring (Ehret and Haack 1981; Ehret 1992), as well as familiar individuals in the context of social recognition (Nelson 1989; Bee and Gerhardt 2001b; this study). Key to such categorizations is the use of decision rules that have been shaped by natural selection. In the context of social recognition based on communication signals, these decision rules should reflect not only the perceptual resolution of a receiver’s sensory system, but also the patterns of individual variation in signals, the payoffs of correct and incorrect decisions, and the rates at which animals are likely to incur these payoffs (Reeve 1989; Wiley 1994). As highlighted by the present study, signal detection theory provides a useful framework for integrating signal analyses, models of receiver utility that consider encounter rates and payoffs, and empirical studies of behavioral discrimination thresholds to better understand both the mechanisms and evolution of decision rules for social recognition.

