## Supplementary Material for "Predicting and measuring decision rules for social recognition in a Neotropical frog"

**Table S-1.** Predicted optimal discrimination thresholds for *temperature-corrected* acoustic properties of calls from data pooled across pulses along with the associated probabilities of different outcomes of correct and incorrect decisions.

| | | | Within-individual differences (%) | | Between-individual differences (%) | |
|---|---|---|---|---|---|---|
| Call Property | Detectability | Predicted threshold (%) | Below threshold ($1 - p_F$) | Above threshold ($p_F$) | Above threshold ($p_D$) | Below threshold ($1 - p_D$) |
| Pulse Duration | 0.24 | 6.6 | 83 | 17 | 60 | 40 |
| Pulse Interval | 0.23 | 9.2 | 81 | 19 | 57 | 43 |
| Pulse Period | 0.28 | 5.1 | 84 | 16 | 63 | 37 |

Probabilities of correct rejections ($1 - p_F$) and false alarms ($p_F$) are computed from the within-individual differences below and above the predicted threshold, respectively. Probabilities of correct detections ($p_D$) and missed detections ($1 - p_D$) are computed from between-individual differences above and below the predicted threshold, respectively.

**Table S-2.** Predicted optimal discrimination thresholds for acoustic properties of calls *without temperature correction* from data pooled across pulses along with the associated probabilities of different outcomes of correct and incorrect decisions.

| | | | Within-individual differences (%) | | Between-individual differences (%) | |
|---|---|---|---|---|---|---|
| Call property | Detectability | Predicted threshold (%) | Below threshold ($1-p_F$) | Above threshold ($p_F$) | Above threshold ($p_D$) | Below threshold ($1 - p_D$) |
| Pulse duration | 0.24 | 7.9 | 85 | 15 | 55 | 45 |
| Pulse interval | 0.27 | 10.4 | 86 | 14 | 57 | 43 |
| Pulse period | 0.32 | 5.4 | 86 | 14 | 66 | 34 |

Probabilities of correct rejections ($1 - p_F$) and false alarms ($p_F$) are computed from the within-individual differences below and above the predicted threshold, respectively. Probabilities of correct detections ($p_D$) and missed detections ($1 - p_D$) are computed from between-individual differences above and below the predicted threshold, respectively.

**Table S-3.** Predicted optimal thresholds for *temperature-corrected* acoustic properties of calls for each pulse analyzed separately along with the associated probabilities of different outcomes of correct and incorrect decisions.

| Call Property | | Detectability | Predicted threshold (%) | Within-individual differences (%) | | Between-individual differences (%) | |
|---|---|---|---|---|---|---|---|
| | | | | Below threshold $(1 - p_F)$ | Above threshold $(p_F)$ | Above threshold $(p_D)$ | Below threshold $(1 - p_D)$ |
| Call Temporal Properties | Call Duration | 0.22 | 4.6 | 63 | 37 | 83 | 17 |
| | Call Interval | 0.19 | 24.1 | 67 | 33 | 63 | 37 |
| | Call Period | 0.18 | 22 | 67 | 33 | 62 | 38 |
| Pulse Duration | Pulse 1 | 0.23 | 6.3 | 79 | 21 | 62 | 38 |
| | Pulse 2 | 0.25 | 6.6 | 85 | 15 | 60 | 40 |
| | Pulse 3 | 0.22 | 6.6 | 82 | 18 | 59 | 41 |
| Pulse Interval | Pulse 1 | 0.25 | 11.5 | 85 | 15 | 56 | 44 |
| | Pulse 2 | 0.22 | 6.7 | 76 | 24 | 60 | 40 |
| Pulse Period | Pulse 1 | 0.28 | 6.4 | 88 | 12 | 59 | 41 |
| | Pulse 2 | 0.28 | 5.1 | 90 | 10 | 59 | 41 |
| Pulse Rise Time | Pulse 1 | 0.13 | 42.2 | 54 | 46 | 66 | 34 |
| | Pulse 2 | 0.11 | 43.1 | 51 | 49 | 65 | 35 |
| | Pulse 3 | 0.10 | 34.6 | 48 | 52 | 69 | 31 |
| Pulse Fall Time | Pulse 1 | 0.11 | 35.6 | 48 | 52 | 70 | 30 |
| | Pulse 2 | 0.10 | 33.4 | 52 | 48 | 65 | 35 |
| | Pulse 3 | 0.12 | 34.5 | 62 | 38 | 58 | 42 |
| Dominant Frequency | Pulse 1 | 0.22 | 1.8 | 71 | 29 | 72 | 28 |
| | Pulse 2 | 0.28 | 1.7 | 79 | 21 | 72 | 28 |
| | Pulse 3 | 0.28 | 1.7 | 81 | 19 | 69 | 31 |

Probabilities of correct rejections $(1 - p_F)$ and false alarms $(p_F)$ are computed from the within-individual differences below and above the predicted threshold, respectively. Probabilities of correct detections $(p_D)$ and missed detections $(1 - p_D)$ are computed from between-individual differences above and below the predicted threshold, respectively. The predicted threshold for each call property was the threshold for discrimination that resulted in the largest difference between the probabilities of correct detection and false alarm.

**Table S-4.** Predicted optimal thresholds for acoustic properties of calls *without temperature correction* for each pulse analyzed separately along with the associated probabilities of different outcomes of correct and incorrect decisions.

| Call Property | | Detectability | Predicted threshold (%) | Within-individual differences (%): Below threshold ($1 - p_F$) | Within-individual differences (%): Above threshold ($p_F$) | Between-individual differences (%): Above threshold ($p_D$) | Between-individual differences (%): Below threshold ($1 - p_D$) |
|---|---|---|---|---|---|---|---|
| Call Temporal Properties | Call Duration | 0.23 | 4.8 | 64 | 36 | 85 | 15 |
| | Call Interval | 0.17 | 19.8 | 66 | 34 | 63 | 37 |
| | Call Period | 0.16 | 19.7 | 68 | 32 | 59 | 41 |
| Pulse Duration | Pulse 1 | 0.24 | 7.9 | 84 | 16 | 56 | 44 |
| | Pulse 2 | 0.26 | 7.9 | 87 | 13 | 56 | 44 |
| | Pulse 3 | 0.23 | 8.4 | 85 | 15 | 53 | 47 |
| Pulse Interval | Pulse 1 | 0.27 | 10.4 | 82 | 18 | 62 | 38 |
| | Pulse 2 | 0.26 | 9 | 88 | 12 | 54 | 46 |
| Pulse Period | Pulse 1 | 0.32 | 6.4 | 88 | 12 | 64 | 36 |
| | Pulse 2 | 0.32 | 5.2 | 90 | 10 | 63 | 37 |
| Pulse Rise Time | Pulse 1 | 0.13 | 41 | 52 | 48 | 67 | 33 |
| | Pulse 2 | 0.11 | 41.7 | 47 | 53 | 68 | 32 |
| | Pulse 3 | 0.10 | 36.9 | 48 | 52 | 69 | 31 |
| Pulse Fall Time | Pulse 1 | 0.12 | 30.5 | 45 | 55 | 73 | 27 |
| | Pulse 2 | 0.11 | 38.1 | 56 | 44 | 61 | 39 |
| | Pulse 3 | 0.12 | 32.2 | 61 | 39 | 59 | 41 |
| Dominant Frequency | Pulse 1 | 0.23 | 2.3 | 71 | 29 | 68 | 32 |
| | Pulse 2 | 0.29 | 2 | 79 | 21 | 67 | 33 |
| | Pulse 3 | 0.28 | 2.3 | 82 | 18 | 63 | 37 |

Probabilities of correct rejections ($1 - p_F$) and false alarms ($p_F$) are computed from the within-individual differences below and above the predicted threshold, respectively. Probabilities of correct detections ($p_D$) and missed detections ($1 - p_D$) are computed from between-individual differences above and below the predicted threshold, respectively. The predicted threshold for each call property was the threshold for discrimination that resulted in the largest difference between the probabilities of correct detection and false alarm.

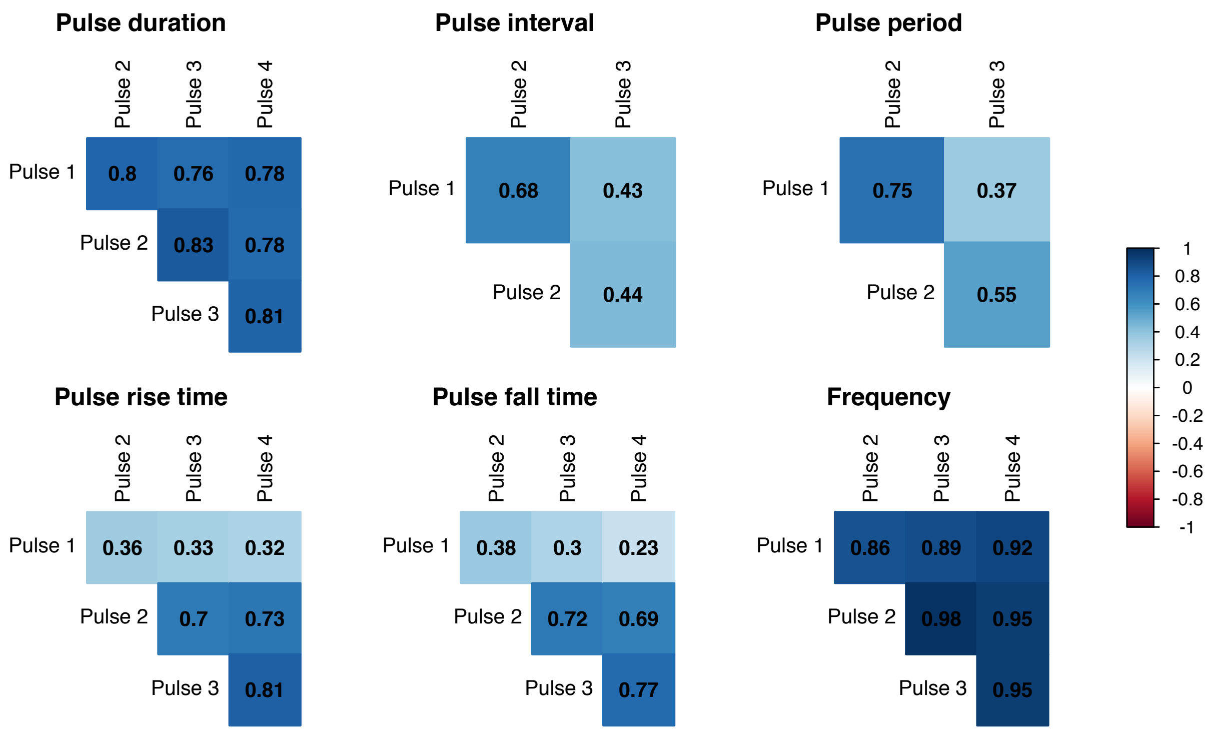


**Figure S-1.** Matrices displaying Pearson correlation coefficients of the correlation between different pulses within calls in acoustic properties. We do not display correlations with pulses five and six because our dataset has only seven and one observations of those pulses, respectively.

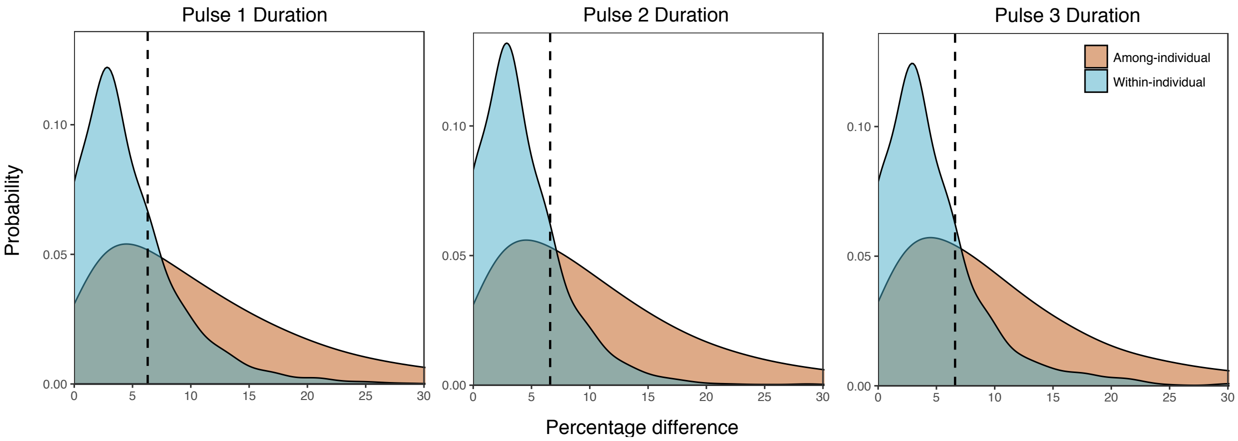


**Figure S-2.** Probability density functions of within-individual and among-individual percentage differences in pulse duration for the three pulses in a call. The dotted line in each graph represents the predicted optimal discrimination threshold for each pulse from an analysis of call variation.

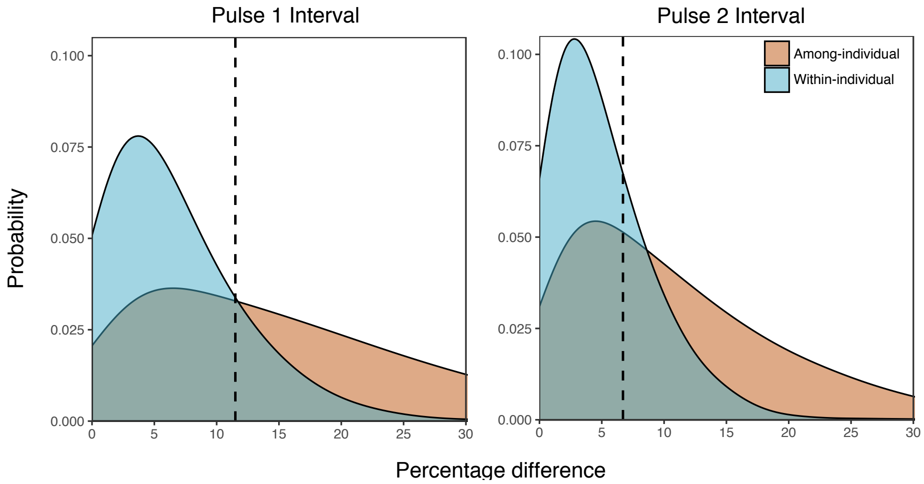


**Figure S-3.** Probability density functions of within-individual and among-individual percentage differences in pulse interval for the two intervals in a call. The dotted line in each graph represents the predicted optimal discrimination threshold for each interval from an analysis of call variation.

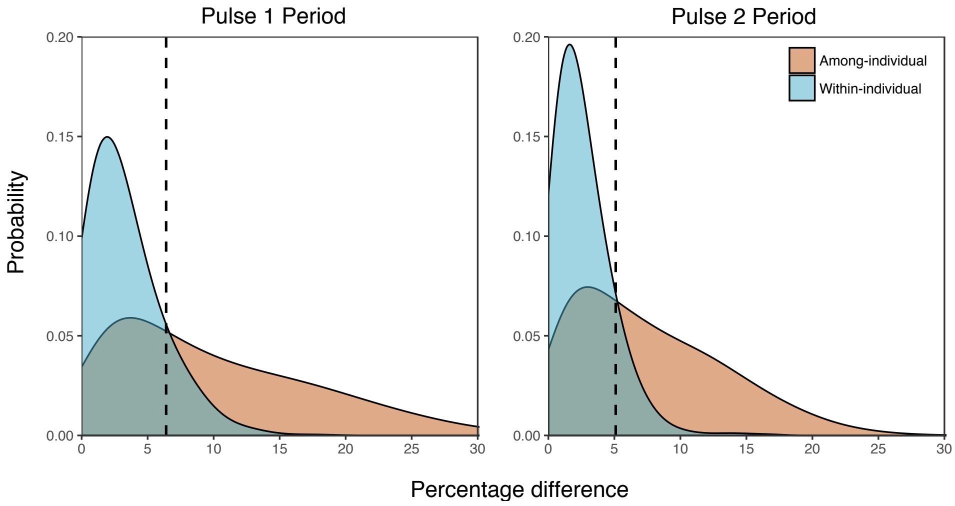


**Figure S-4.** Probability density functions of within-individual and among-individual percentage differences in pulse periods for the two pulse periods in a call. The dotted line in each graph represents the predicted optimal discrimination threshold for each pulse period from an analysis of call variation.

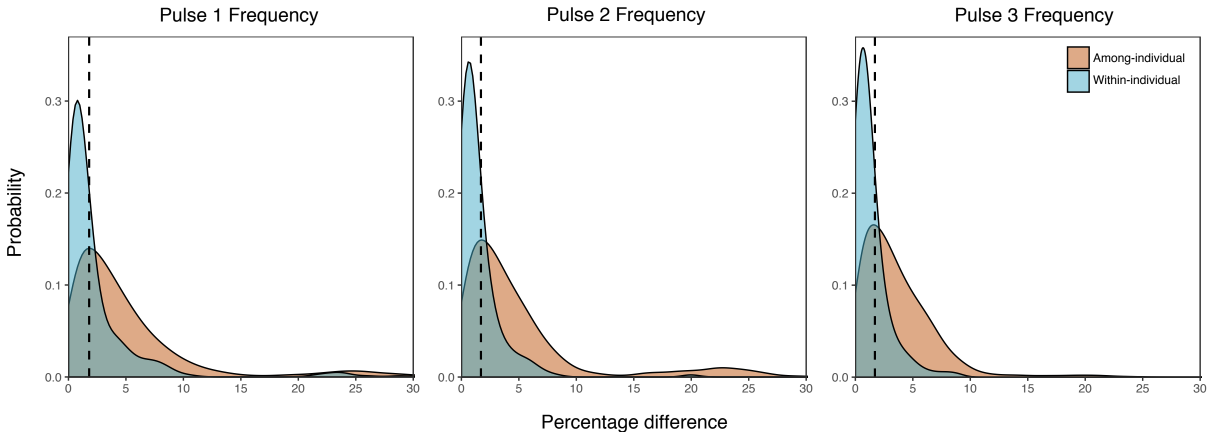


**Figure S-5.** Probability density functions of within-individual and among-individual percentage differences in dominant frequency of the three pulses in a call. The dotted line in each graph represents the predicted optimal discrimination threshold for each pulse from an analysis of call variation.

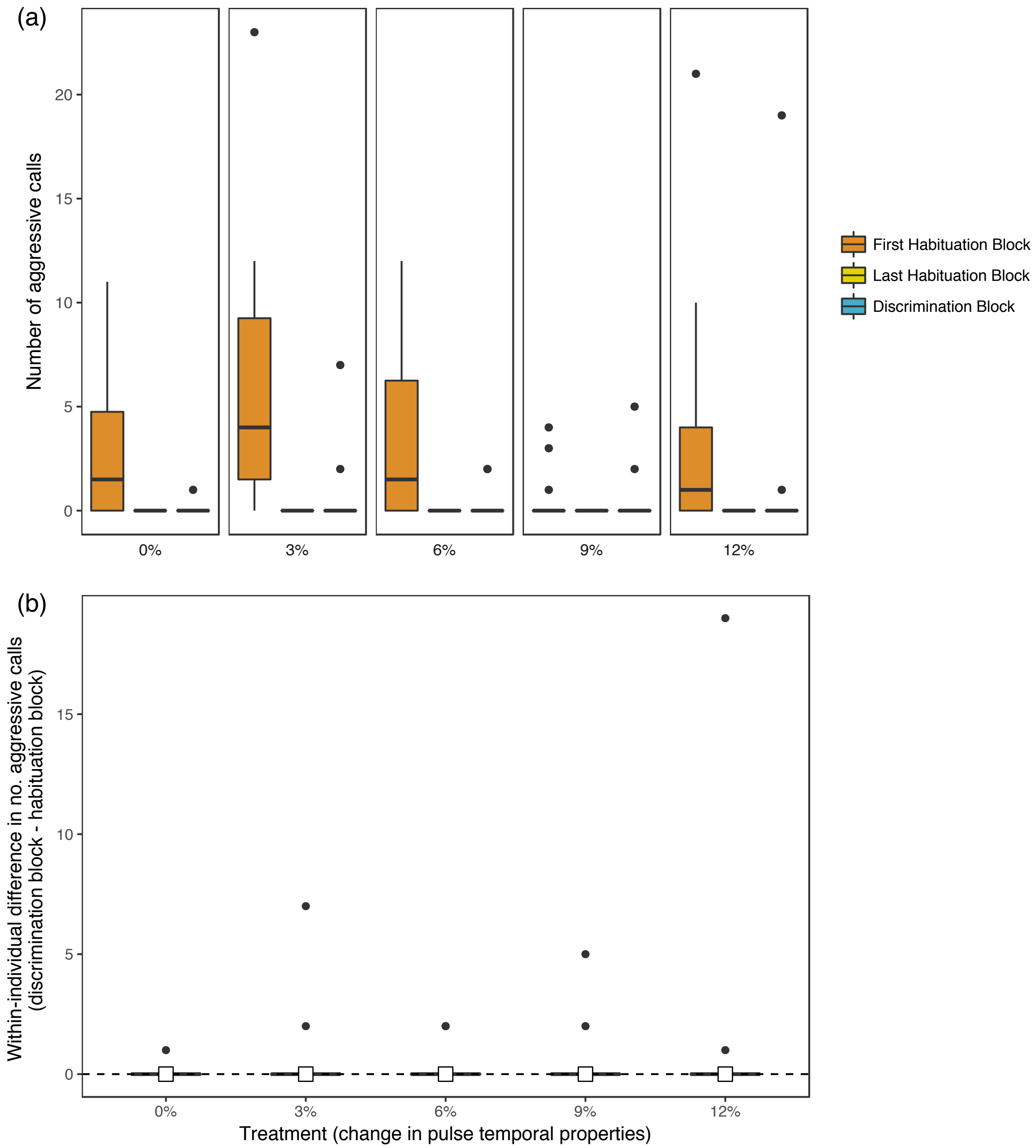


**Figure S-6.** (a) Boxplots of the number of aggressive calls produced by territorial male subjects during the first and last blocks of the habituation phase and during the block of the discrimination phase for each treatment. (b) Boxplots showing the within-individual differences in number of aggressive calls between the last block of the habituation phase and the discrimination phase for each treatment. Horizontal bars represent the median, box hinges represent the interquartile range, whiskers extend to the range but no further than 1.5 times the interquartile range, and outliers are shown as points. White squares also show the median in (b) to aid visualization.

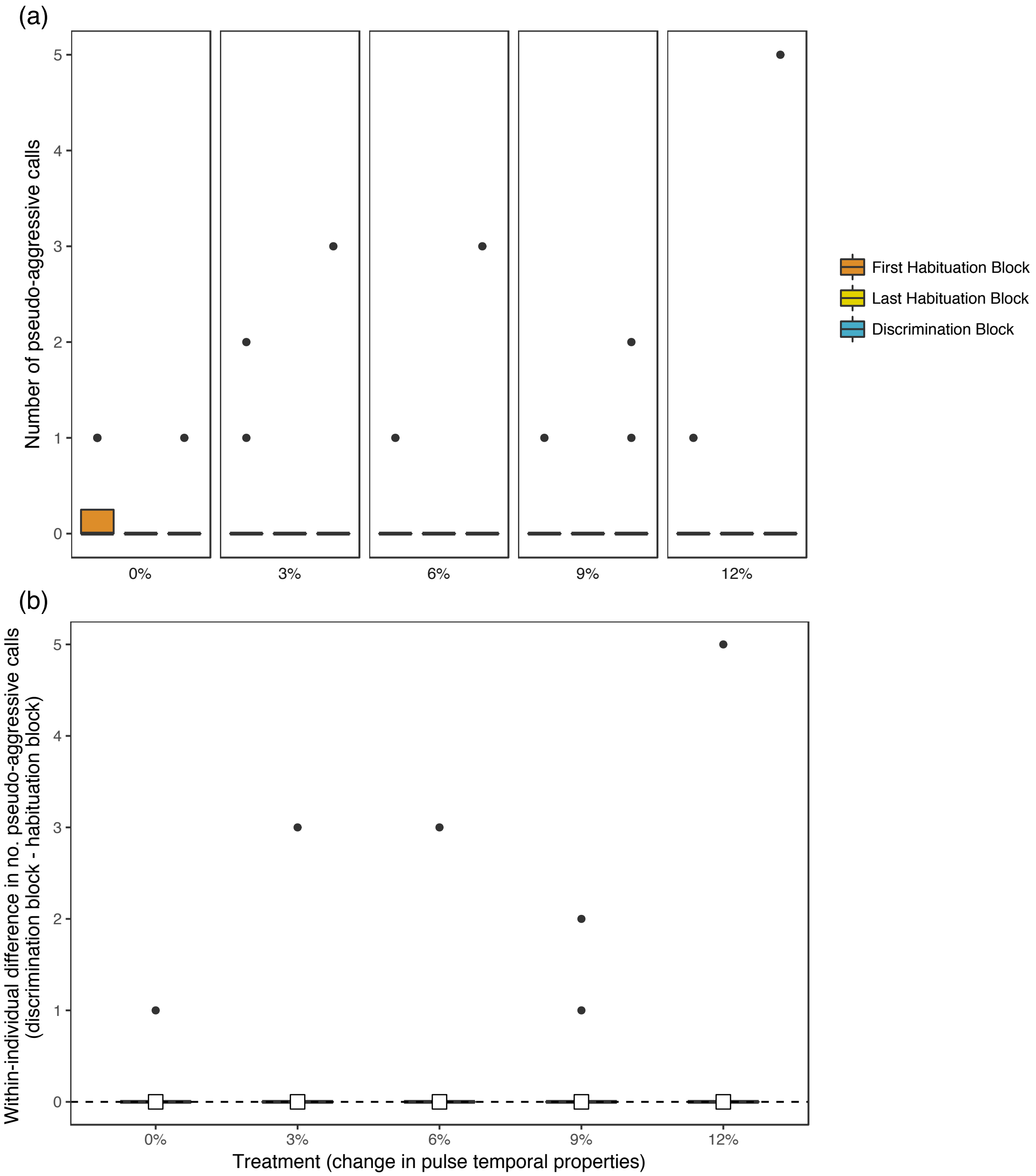


**Figure S-7.** (a) Boxplots of the number of pseudo-aggressive calls produced by territorial male subjects during the first and last blocks of the habituation phase and during the block of the discrimination phase for each treatment. (b) Boxplots showing the within-individual differences in number of pseudo-aggressive calls between the last block of the habituation phase and the discrimination phase for each treatment. Horizontal bars represent the median, box hinges represent the interquartile range, whiskers extend to the range but no further than 1.5 times the interquartile range, and outliers are shown as points. White squares also show the median in (b) to aid visualization.

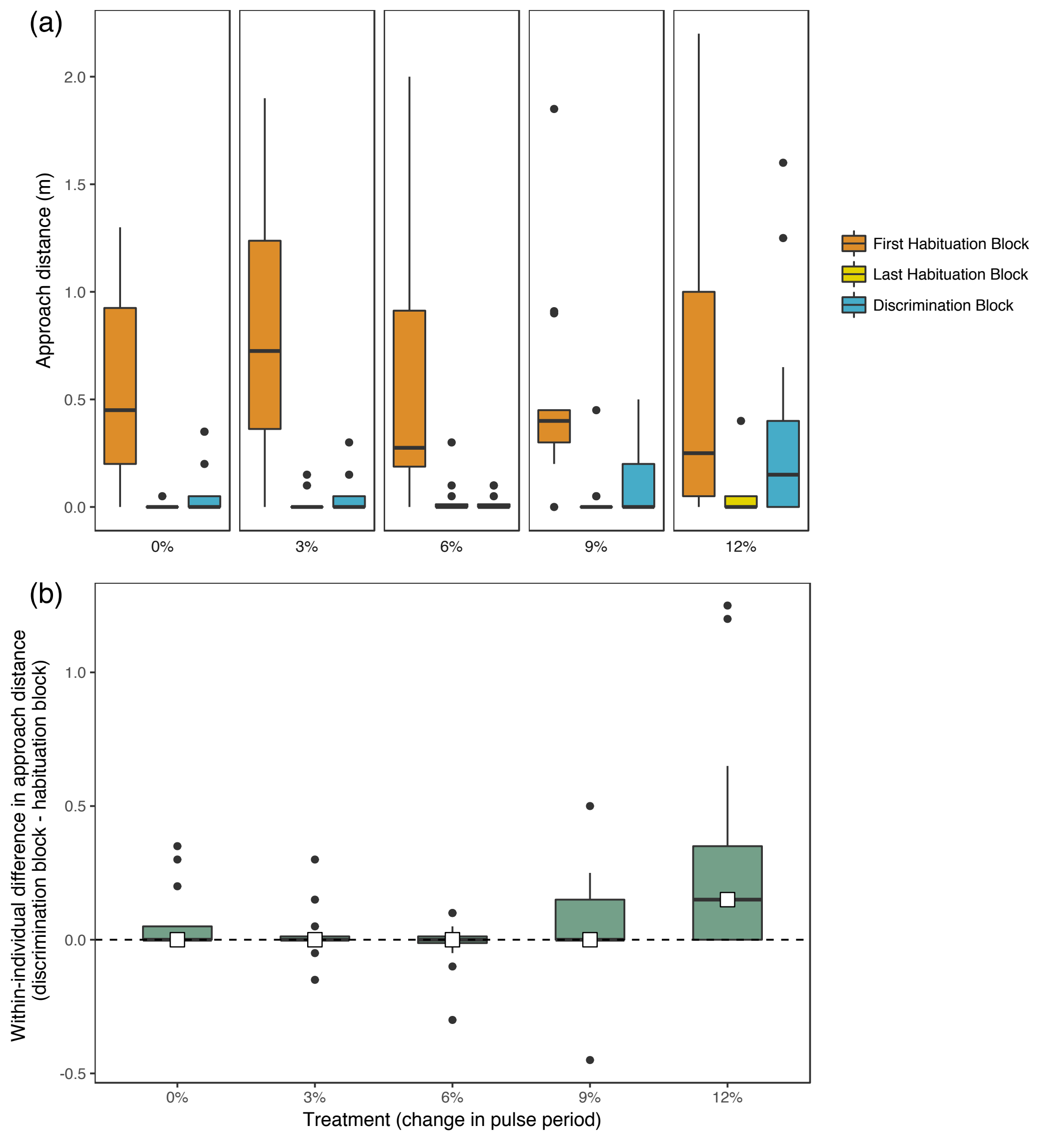


**Figure S-8.** (a) Boxplots of the approach distance of territorial male subjects during the first and last blocks of the habituation phase and during the block of the discrimination phase for each treatment. (b) Boxplots showing the within-individual differences in approach distance between the last block of the habituation phase and the discrimination phase for each treatment. Horizontal bars represent the median, box hinges represent the interquartile range, whiskers extend to the range but no further than 1.5 times the interquartile range, and outliers are shown as points. White squares also show the median in (b) to aid visualization.

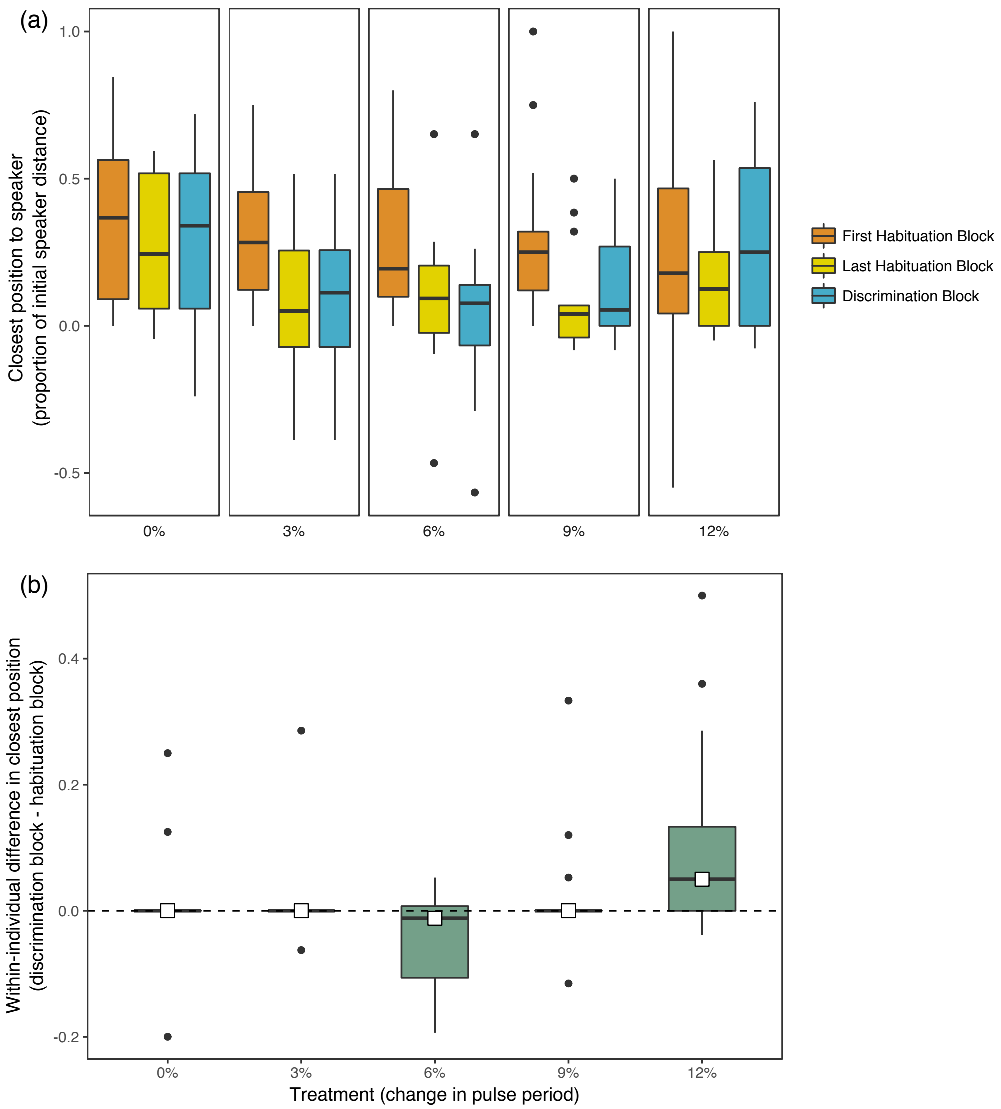


**Figure S-9.** (a) Boxplots of the closest position to the speaker of territorial male subjects, expressed as a proportion of the initial speaker distance, during the first and last blocks of the habituation phase and during the block of the discrimination phase for each treatment. (b) Boxplots showing the within-individual differences in closest position to the speaker between the last block of the habituation phase and the discrimination phase for each treatment. Horizontal bars represent the median, box hinges represent the interquartile range, whiskers extend to the range but no further than 1.5 times the interquartile range, and outliers are shown as points. White squares also show the median in (b) to aid visualization.
